# multiTEMPTED: Joint Dimensionality Reduction of Longitudinal Multi-omic Data with Modality-Specific Temporal Dynamics

**DOI:** 10.64898/2026.09.01.748608

**Authors:** Lou Lindsley, Annabel Settle, Pixu Shi

## Abstract

Longitudinal multi-omic studies profile multiple molecular layers, such as microbiome composition, metabolomics, lipidomics, and proteomics, repeatedly over time. These layers reflect shared subject-level biological processes yet each may exhibit its own temporal dynamics. Most existing methods either integrate multiple omics modalities cross-sectionally or model a single modality longitudinally. The few methods that handle longitudinal multi-omic data assume a shared temporal trajectory across all modalities, limiting their ability to capture modality-specific dynamics, and can be computationally prohibitive at the scale of modern cohort studies. We introduce multiTEMPTED, an extension of the temporal tensor decomposition framework to *M* ≥ 1 simultaneous omic modalities. The method jointly estimates subject components shared across modalities, modality-specific feature loadings identifying the contributing molecular features, and modality-specific temporal trajectories, while accommodating arbitrary, unaligned sampling schedules across subjects and modalities without imputation. In the MOMS-PI pregnancy cohort, joint analysis of the vaginal microbiome and cervicovaginal cytokines yields clearer separation between preterm and term birth outcomes than microbiome data alone, implicates *Lactobacillus*-dominant communities as protective, and identifies diverging cytokine trajectories in women who subsequently deliver preterm. In an exercise study profiling four plasma omics modalities, the leading unsupervised subject component from multiTEMPTED is strongly associated with sex, whereas the leading components from MEFISTO show no discernible association with known study phenotypes and appear numerically degenerate. In simulation experiments, multiTEMPTED accurately recovers the underlying component structure across a range of noise levels, outperforms MEFISTO in multiple settings, and runs several orders of magnitude faster. multiTEMPTED is implemented in the open-source R package multi.tempted (https://github.com/loulind/multi.tempted).

## 1 Introduction

Advances in high-throughput sequencing and mass spectrometry now make it practical to profile multiple molecular layers from the same individuals at multiple time points. Longitudinal multi-omic data, spanning layers such as the microbiome, metabolome, lipidome, proteome, and transcriptome, capture the dynamic interplay among molecular systems and can reveal mechanisms underlying disease progression, host-microbe interactions, and responses to environmental or therapeutic perturbations that cross-sectional snapshots cannot uncover [1]. A growing number of large-scale cohort studies now collect such data, including longitudinal profiling of the vaginal microbiome and host immune markers during pregnancy [2], plasma multi-omic responses to a controlled physiological challenge [3], gut microbiome and metabolome dynamics during dietary fiber intervention and antibiotic recovery [4], gut microbial and metabolomic trajectories in inflammatory bowel disease [5], and multi-omics of host-microbe dynamics across the prediabetes transition [6].

Despite this growth, dedicated analytical methods have lagged behind. Widely used multiomic integration methods, including MOFA [7], MOFA+ [8], and JIVE [9], are designed for cross-sectional data and do not model temporal structure. Methods developed specifically for longitudinal single-omic data, such as CTF [10], microTensor [11], and TEMPTED [12], capture temporal variation but are limited to a single modality. MEFISTO [13], an extension of MOFA with Gaussian process priors over time, is, to the best of our knowledge, the only method targeting continuous longitudinal multi-omic data. However, MEFISTO enforces a single temporal trajectory shared across all modalities within each latent component, an assumption that may not hold when different molecular layers respond to the same biological process with distinct dynamics. Furthermore, the Gaussian process formulation underlying MEFISTO incurs substantial computational cost that scales poorly with the number of subjects and time points, limiting its practicality for large cohort studies.

Jointly analyzing high-dimensional, irregularly sampled, multi-modal longitudinal data introduces further statistical challenges. Individual molecular layers are typically high-dimensional, with far more features than subjects, requiring both regularization to recover interpretable low-dimensional structure and efficient computation to remain tractable at scale. Sampling schedules in cohort studies are often irregular and may differ across subjects or across modalities, restricting the class of methods that can be applied without imputing missing observations or forcing all modalities onto a shared time grid.

We introduce multiTEMPTED, an extension of the single-modality temporal tensor decomposition (TEMPTED) framework [12] to *M* ≥ 1 simultaneous omics modalities. The method organizes each modality’s measurements into a temporal tensor and jointly decomposes these tensors into subject components shared across modalities, modality-specific feature loadings identifying the contributing molecular features, and modality-specific temporal loadings estimated nonparametrically, while accommodating arbitrary and unaligned sampling schedules across subjects and modalities without imputation. We apply multiTEMPTED to two longitudinal multi-omic cohort studies: the MOMS-PI pregnancy cohort [2], where jointly profiling the vaginal microbiome and cervicovaginal cytokines reveals biological structure that microbiome data alone cannot capture; and a plasma multi-omic exercise challenge [3], where multiTEMPTED runs orders of magnitude faster than MEFISTO and yields clearer subject-level separation across four plasma omics modalities. We validate multiTEMPTED through simulation experiments, demonstrating accurate recovery of the latent component structure, with substantially better accuracy and speed than MEFISTO in all settings evaluated.

## 2 Materials and Methods

We model longitudinal multi-omic data as a collection of *M* third-order temporal tensors, one per modality, with modes indexing subjects, molecular features, and time. The tensors are linked across modalities through shared subject loadings, reflecting the premise that the same biological differences between subjects drive coordinated signals across molecular layers. The formulation generalizes the single-modality TEMPTED model [12] by pooling information across modalities when estimating the subject representation, while allowing each modality to carry its own feature loadings and temporal dynamics.

Let *m* = 1, …, *M* denote modality, *i* = 1, …, *n* denote subjects, *j* = 1, …, *p*^(*m*)^ denote features in modality *m*, and 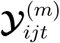 denote the observed value of feature *j* of modality *m* from subject *i* at time points 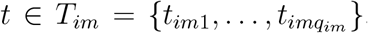. Here, *T*_*im*_ ⊂ *T* is the set of *q*_*im*_ observed time points for subject *i* in modality *m*. Both the count *q*_*im*_ and the observation times themselves may differ across subjects and modalities, accommodating varying temporal sampling and missing time points. 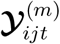 forms an order-3 temporal tensor ***Y***^(*m*)^, with three modes representing subject, feature and time, respectively. We assume the following model for the observed data tensors:

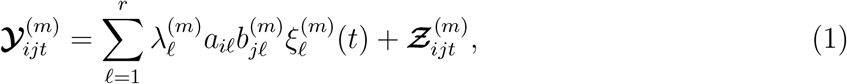

where *r* is the number of low-rank components to approximate the data tensors, *a*_*ℓ*_ = (*a*_1*ℓ*_, …, *a*_*nℓ*_) are subject loadings that are shared across modalities, 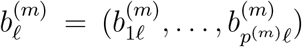 are modality-specific feature loadings quantifying the contribution of each feature to each component, 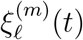 is the modality-specific temporal loading, a single latent function of time common to all subjects and features in modality *m* that characterizes the temporal dynamics of component *ℓ*, and 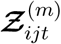 quantifies unexplained remainders and measurement errors. The shared *a*_*ℓ*_ across modalities enable features from multiple modalities to contribute to the same low-dimensional representation of subjects, while the modality-specific 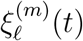 can capture trends that are shared within each modality but differ between modalities. To ensure the identifiability of the model, we require 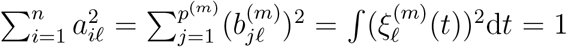 for each *ℓ* = 1, …, *r* and *m* = 1, …, *M*.

The goal of multiTEMPTED is to obtain estimates of 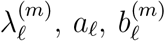 and 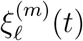 in the approximately CP low-rank structure (1). Denote 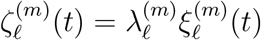. We consider sequentially estimating each component *ℓ* = 1, …, *r* through the following objective function:

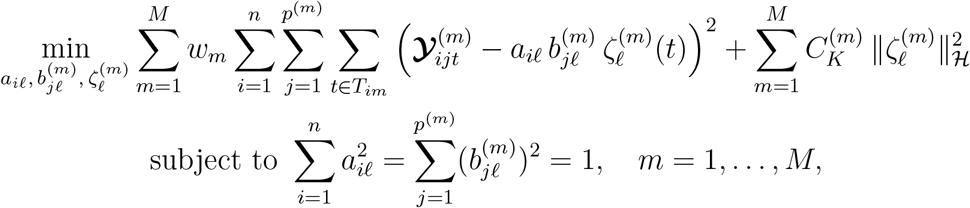

where *w*_1_, …, *w*_*M*_ are weights that adjust different levels of focus on different modalities, and 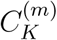 is the modality-specific kernel ridge penalty parameter. The penalty 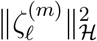 is the squared norm in a reproducing kernel Hilbert space of smooth functions, with 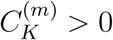 controlling the degree ℋ of regularization, encouraging smooth temporal trajectories and rendering the problem well-posed for irregularly sampled data. Details on the kernel construction and selection of 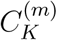 are given in [12]. After obtaining the estimated loadings in component *ℓ* as 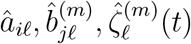, we update ***Y***^(*m*)^ for each *m* = 1, …, *M* by subtracting the previously estimated components using 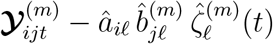. The details of this iterative algorithm are described in Section A.

## 3 Results

### 3.1 MOMS-PI Study

The Multi-Omic Microbiome Study: Pregnancy Initiative (MOMS-PI) [2] enrolled 1,572 pregnant women with repeated sampling throughout gestation. The cohort was enriched for African-American women, a population at elevated risk for spontaneous preterm birth (PTB, delivery before 37 weeks). Two molecular data layers were assessed at each visit: the vaginal microbiome profiled by 16S rRNA gene sequencing and a panel of cytokines measured from cervicovaginal fluid. We applied multiTEMPTED jointly to both modalities and, as a comparison, to the microbiome data alone, in each case fitting *r* = 3 components. Detailed description of the data and preprocessing procedure can be found in Section B.1.

The joint multiTEMPTED embedding shows a clear cluster of subjects containing all PTB subjects in the space of the top two components, compared to no obvious pattern in the microbiome-only embedding, demonstrating that incorporating cytokine dynamics alongside microbial composition yields clearer separation of clinically relevant subject groups (Figure 1a). The microbiome feature loadings on Component 1 highlight the dominant role of *Lactobacillus* species, particularly *L. crispatus* and *L. iners*, consistent with the established protective role of *Lactobacillus*-dominant communities against adverse birth outcomes. Component 2 captures variation attributable to anaerobic taxa such as *Gardnerella* and *Prevotella*, which are associated with bacterial vaginosis and preterm birth risk (Figure 1b). IL-1ra and GM-CSF are the top two cytokine features by loading magnitude in all three components, with consistently opposite signs, making them natural candidates to examine. The temporal loading curves for these cytokines reveal a sustained separation between PTB and term pregnancies throughout gestation: women who deliver preterm exhibit consistently higher IL-1ra and consistently lower GM-CSF than those who deliver at term (Figure 1c). These patterns are consistent with a model of premature immune activation and loss of anti-inflammatory tone preceding spontaneous preterm delivery [14, 15].

**Figure 1:**
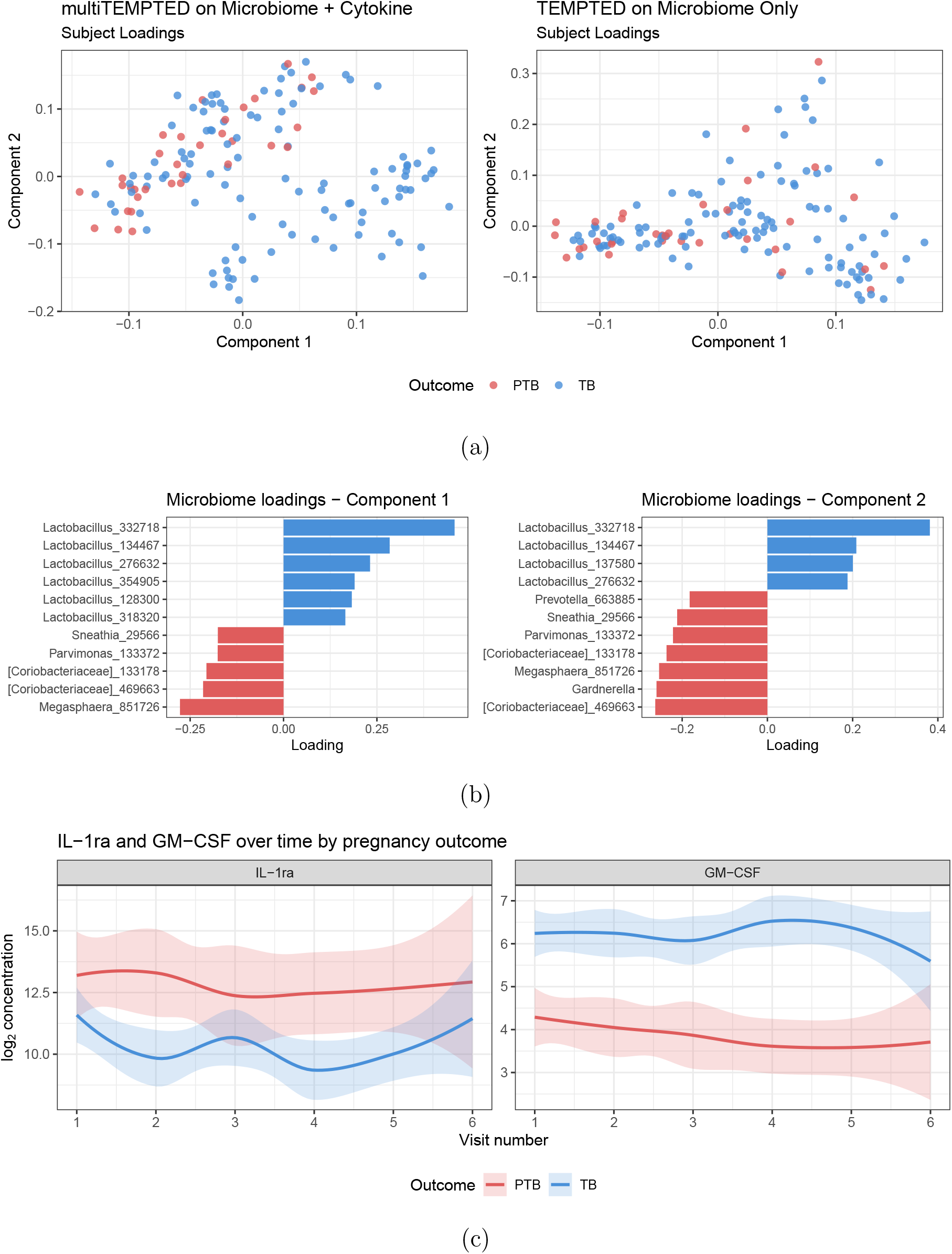
MOMS-PI study. (a) First two subject components from joint microbiome + cytokine analysis and microbiome-only analysis, coloured by pregnancy outcome (PTB: preterm birth; TB: term birth). (b) Top microbiome feature loadings for components 1 and 2 from the joint multiTEMPTED run. (c) Smoothed mean trajectories (LOESS*±*SE) of IL-1ra and GM-CSF concentrations over six visits, stratified by pregnancy outcome.

### 3.2 iPOP Exercise Study

The iPOP Exercise Study, detailed in [3], profiled 36 healthy volunteers with four plasma omics modalities: untargeted metabolomics (LC-MS), semi-targeted lipidomics (Lipidyzer), targeted proteomics (immunoassays), and untargeted proteomics (SWATH-MS). All four assays were drawn from the same plasma sample, taken at five time points spanning a single controlled session: at rest before exercise, and at 2, 15, and 30 minutes and 1 hour after a symptom-limited cardiopulmonary exercise test performed following an overnight fast. After preprocessing (Appendix B.2), the modalities contribute 674 metabolites, 122 aggregated lipid features, and 369 protein markers (targeted and untargeted proteomics concatenated into a single modality).

We compare multiTEMPTED with MEFISTO [13], the temporal extension of MOFA [7] and, to our knowledge, the only other method targeting low-rank structure in continuous longitudinal multi-omic data. We applied both methods to the iPOP data with *r* = 3 components, using equal modality weights. Since MEFISTO requires a group label per subject, to run dimensionality reduction without group-label supervision, we assigned each subject to its own group. Sex was not provided to either method during fitting and serves as an external validation variable. multiTEMPTED’s leading subject component is markedly more strongly associated with sex than any MEFISTO factor (maximum |cor| of 0.53 against 0.16; Figure 2), whereas MEFISTO’s factor loadings appear numerically degenerate, with the two sexes intermixed throughout the factor space.

**Figure 2:**
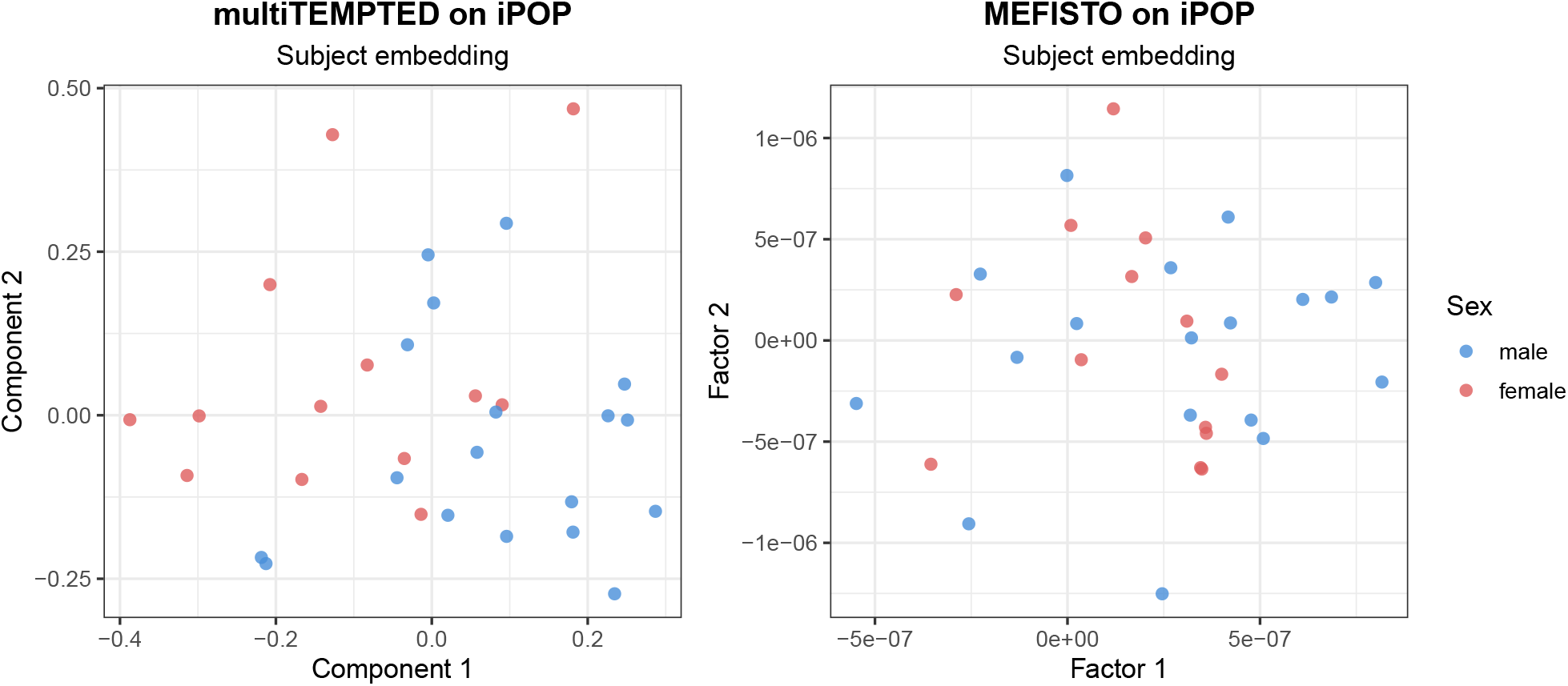
iPOP study. First two subject components from each method, coloured by sex. multiTEMPTED’s embedding separates the two groups more cleanly, despite sex never entering either fit.

### 3.3 Synthetic Data Simulation

We conducted three simulation experiments in a progressive sequence. The first evaluates whether multiTEMPTED accurately recovers the true underlying signal under its own model. The second tests multiTEMPTED against MEFISTO when each modality follows its own temporal pattern. The third tests whether multiTEMPTED remains competitive when all modalities share the same temporal dynamics, as MEFISTO assumes. In Simulations 2 and 3, MEFISTO is run at its recommended configuration. Since MEFISTO outputs each factor as a subject-by-time matrix rather than separate subject and temporal loadings, we extract subject and temporal loadings via rank-one SVD to enable direct comparison. Full simulation details are given in Appendix C.

#### Simulation 1: recovery of the underlying signal across irregularly sampled modalities

The first experiment evaluates whether multiTEMPTED accurately recovers the true underlying signal at a wide range of noise levels when each of the three modalities is given a distinct temporal pattern. Each subject and modality has its own independently drawn measurement times. At low noise, multiTEMPTED accurately recovered all temporal curves across all three modalities and three components (Figure 3, left). Subject scores, feature contributions, and temporal trajectories remained accurate until noise became very large and degraded smoothly in parallel rather than collapsing abruptly (Figure 3, right). Recovery was unaffected by the complete lack of shared time points across subjects and modalities.

**Figure 3:**
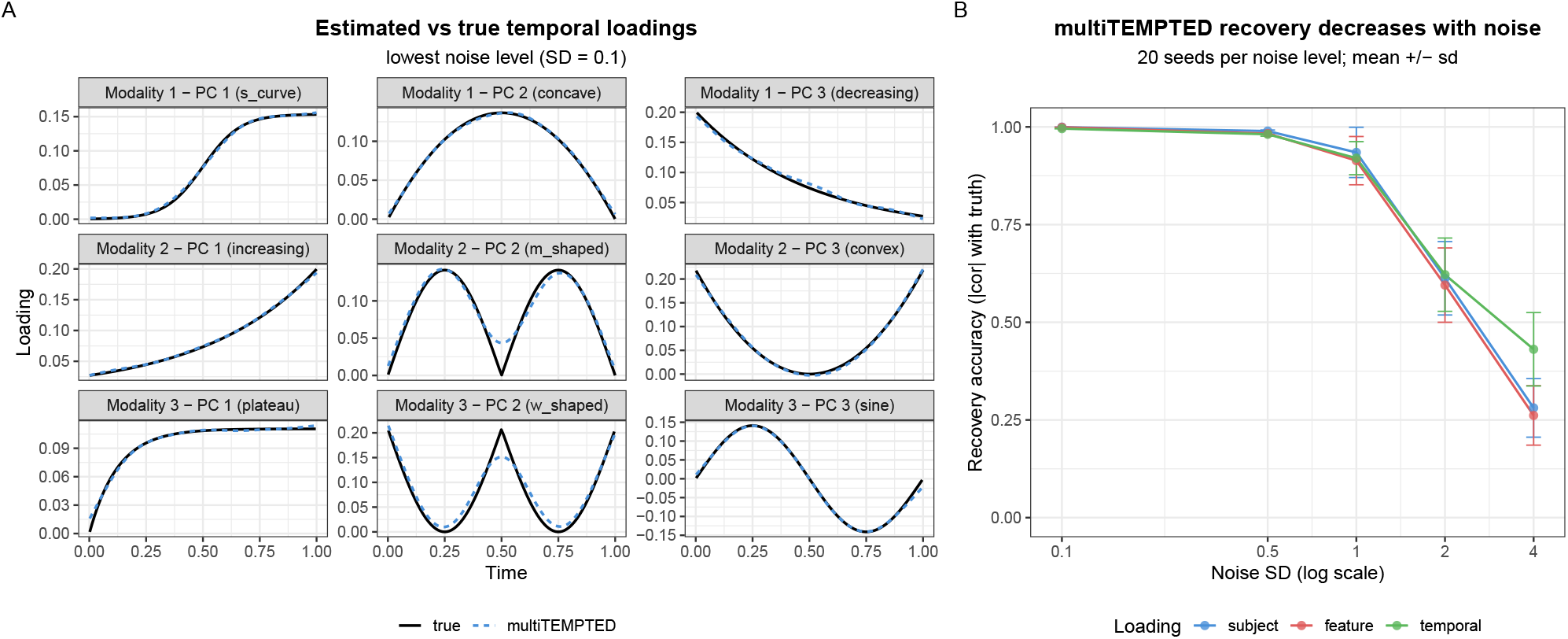
Simulation 1. Recovery accuracy against noise level, and estimated versus true temporal loadings at the lowest noise level. multiTEMPTED recovers the time trajectories despite sampling times being unaligned across both modality and subject.

#### Simulation 2: each modality follows its own temporal pattern

The second experiment simulates three modalities with each having its own temporal pattern within each component, reflecting the scenario where one modality may respond to a biological stimulus faster than others, even when all reflect the same underlying biology. At a low noise level, MEFISTO, which fits a single shared temporal pattern per component, produced the same curve for all three modalities within a given factor, demonstrating its inability to capture the modality-specific dynamics. In contrast, multiTEMPTED correctly recovered the temporal pattern for each modality in each component on every simulation run (Figure 4). Computationally, multiTEMPTED completed each run in under one second, while MEFISTO required several hours per run.

**Figure 4:**
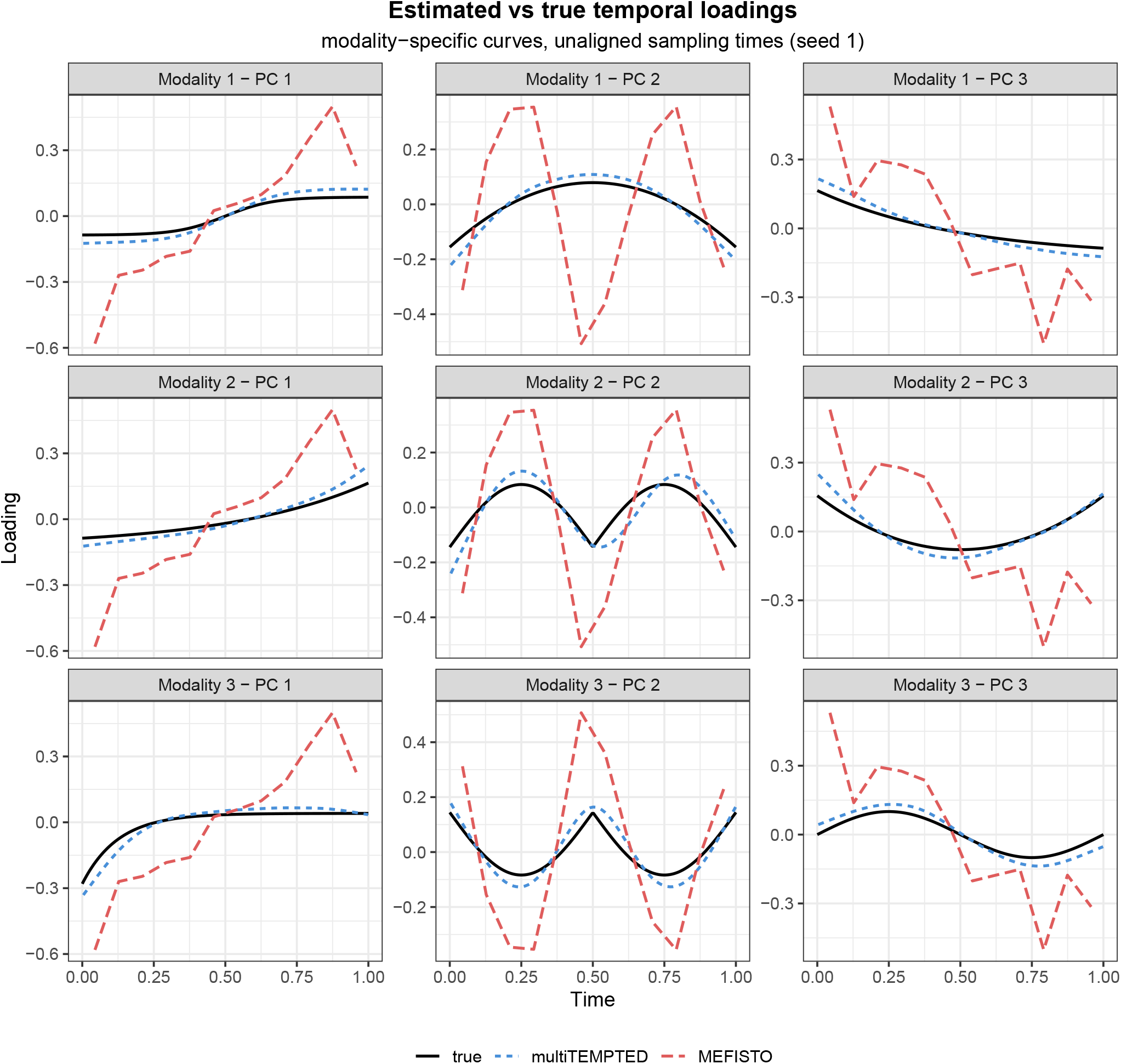
Simulation 2. Estimated versus true temporal patterns for each (modality, component) pair on a representative run. multiTEMPTED tracks each modality’s distinct curve; MEFISTO fits one trajectory per component shared across all modalities and cannot match them simultaneously.

#### Simulation 3: multiTEMPTED under the dynamics MEFISTO assumes

The third experiment steps back to a scenario more favorable to MEFISTO: all three modalities share the same temporal pattern within each component, and subjects fall into distinct biological groups. Sampling followed a regular schedule with per-subject jitter and all modalities co-measured at each visit (details in Appendix C). At a moderate noise level, MEFISTO misclassified between 20% and 60% of subjects across runs, with a median near 47%, while multiTEMPTED correctly classified subjects in all 100 simulation runs (Figure 5, left). Recovery of the temporal patterns showed the same ordering: multiTEMPTED achieved superior temporal dynamic recovery, with a median |cor| = 0.97 and a tight spread compared to MEFISTO’s highly varied |cor| between 0.50 and 0.94 (Figure 5, centre). multiTEMPTED completed all 100 runs in approximately 11 seconds, while MEFISTO required 3.6 days in total (for elapsed computation time per run, see Figure 5, right).

**Figure 5:**
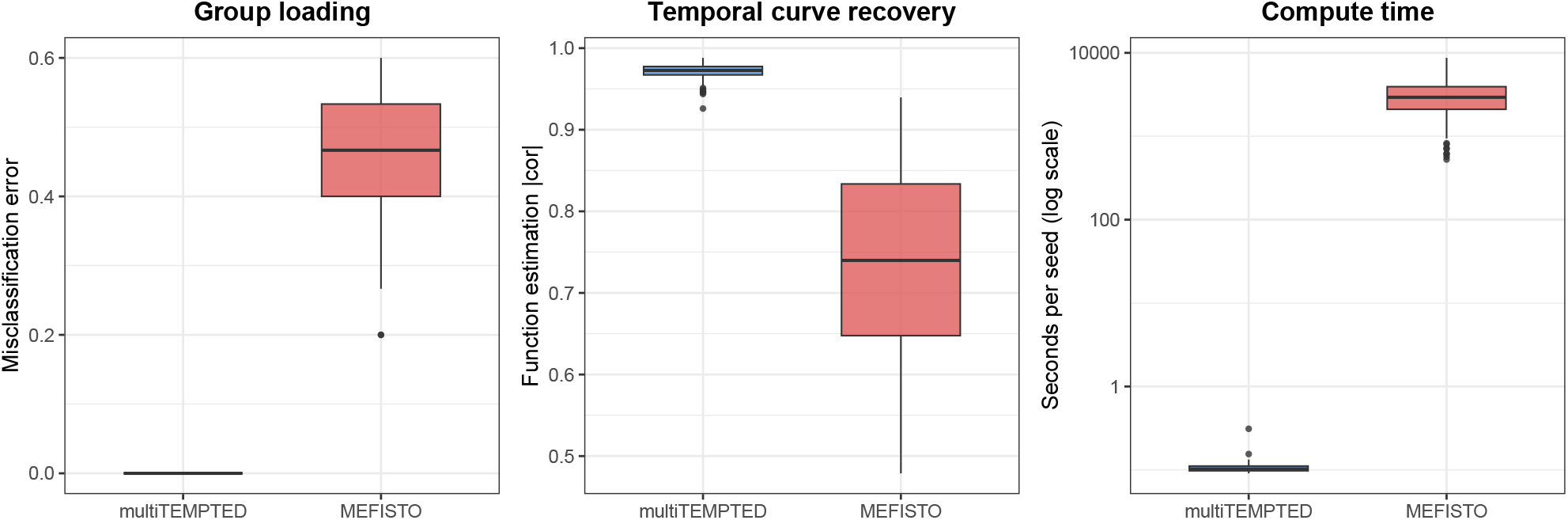
Simulation 3. Distribution over 100 simulation runs of subject-group misclassification error (left), temporal loading recovery (centre), and elapsed computation time per run measured by `system.time()` in R on a logarithmic scale (right). Even when all modalities share the same temporal dynamics, multiTEMPTED identifies subject groups without error and is several orders of magnitude faster than MEFISTO.

## 4 Discussion

We have introduced multiTEMPTED, a tensor decomposition framework for dimensionality reduction of longitudinal multi-omic data. By sharing subject loadings across modalities while allowing each modality its own nonparametric temporal trajectory, the method captures coordinated biological variation without imposing a common temporal model. In simulation experiments, multiTEMPTED accurately recovered the latent component structure, outperformed MEFISTO under both modality-specific and shared temporal dynamics, and ran several orders of magnitude faster. Applied to the MOMS-PI pregnancy cohort, the joint microbiome-cytokine analysis revealed a preterm birth signature that microbiome data alone could not provide, with diverging IL-1ra and GM-CSF trajectories emerging as robust signals. In the iPOP exercise study, multiTEMPTED yielded a clear sex-associated subject embedding across four plasma omics modalities while MEFISTO produced numerically degenerate solutions on the same data.

Beyond its empirical performance, multiTEMPTED has several practical properties worth noting. A particularly useful one is that components are estimated sequentially, with deflation after each step. This means the quality of any given component is unaffected by the choice of total rank *r*: earlier components are locked in before later ones are extracted, so a practitioner can examine the leading components without committing to a final rank upfront and add more components incrementally as needed. In addition to this flexibility, multiTEMPTED is designed as an unsupervised dimensionality reduction tool. It returns point estimates of subject loadings, feature loadings, and temporal curves rather than inferential quantities. Because it is unsupervised, the method surfaces the dominant axes of variation in the data without any guarantee that those axes will align with a specific phenotype of interest. Downstream inference about biological associations, such as relating subject scores to phenotypes, is naturally carried out with standard statistical tests applied to the extracted embeddings.

One area for future methodological development is modality weighting. The framework supports modality-specific weights, but all analyses here used equal weights. Principled guidance on weight selection is lacking, and equal weighting may not be appropriate when modalities differ substantially in feature dimensionality, noise level, or biological relevance to the question at hand. Adaptive or data-driven weighting schemes that balance the influence of each modality could improve performance in heterogeneous settings and would be a natural extension of the framework. Other directions include a supervised or semi-supervised variant that incorporates phenotype information to guide component extraction toward biologically relevant axes of variation, and extension to data types with non-Gaussian structure, such as count-based or compositional measurements, which the current implementation handles only after preprocessing to approximate normality.

## Appendix

### A Estimation Algorithm

The sequential estimation procedure ensures the uniqueness of each component regardless of the chosen rank *r* and allows users to explore additional components without impacting those previously obtained. The updates of 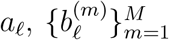 and 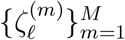 can be obtained iteratively without enforcing the unit-norm constraints, and the modality-specific scales 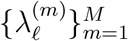 can then be obtained through one step of linear regression after the loadings reach convergence and are normalized to norm one.

For each modality *m*, set 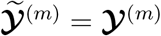 or ***Y***^(*m*)^ after mean subtraction. For each component *ℓ* = 1, …, *r* sequentially, we perform the following Steps 1 to 3 to estimate 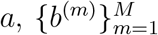 and 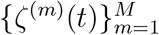. (Throughout Steps 1–3 we suppress the component index *ℓ* on the loadings to lighten notation.)

Step 1: (Initialization) Initialize the shared subject loading as 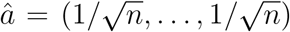. For each modality *m* = 1, …, *M*, set 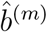 as the first left singular vector of mode-2 matricization of 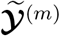: i.e., the 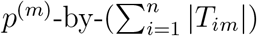 matrix with 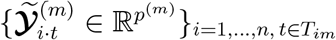 as its columns.

Step 2: (Estimation of loadings) To estimate the loadings, we minimize the following function by iteratively updating the modality-specific temporal loadings 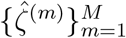, the shared subject loading *â*, and the modality-specific feature loadings 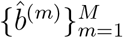 until convergence:

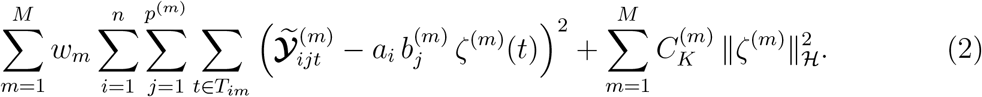

The penalty 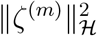 encourages smooth temporal trajectories and renders the problem well-posed for irregularly sampled data. See Section 2 for details on the kernel construction and selection of 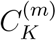.

a. For each *m* = 1, …, *M*, update 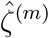 separately by applying kernel ridge regression to solve

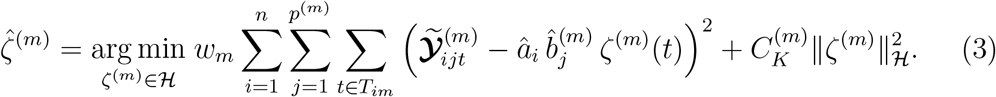

The details of this update are similar to the single modality version in [12].
b. Jointly update the shared subject loading *â* = (*â*_1_, …, *â*_*n*_) across all modalities by

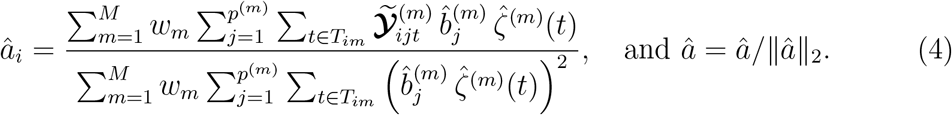
c. For each *m* = 1, …, *M*, update 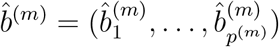 separately by

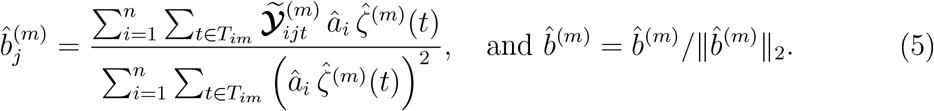

(a)-(c) are iterated until convergence, defined as the *ℓ*_2_ norm of the changes in the subject loading *â* and each feature loading 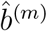 falling below 10^−4^.

Step 3: (Subtracting previous components) For each *m* = 1, …, *M*, normalize 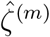 to 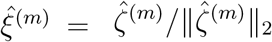 and update 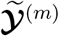 by

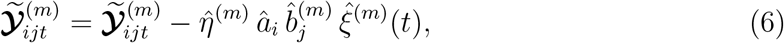

where 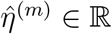 is obtained by solving the following least squares problem:

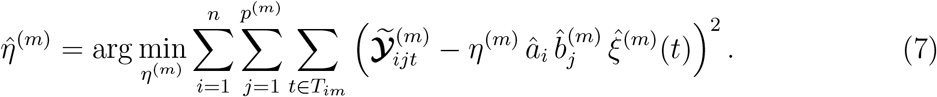

Step 4: After obtaining 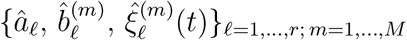 by sequentially running Steps 1–3, we estimate the modality-specific scales 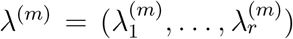 for each *m* = 1, …, *M* via the following least squares problem:

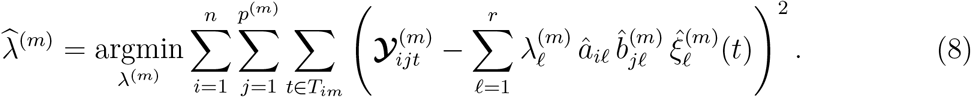

### B Preprocessing of Case Studies

#### B.1 MOMS-PI Study

The 16S microbiome data, cytokine data and associated metadata from the MOMS-PI study was downloaded from the GitHub repository of R package HMP2Data https://github.com/dozmorovlab/HMP2Data/. The pregnancy outcome data was downloaded from the GitHub repository https://github.com/pixushi/TEMPTED_paper. From the full cohort, we focused on the first six time points where majority of the subjects were measured, and analyzed a subset of 34 women with spontaneous PTB and 111 term birth controls with at least two microbiome measurements and two cytokine measurements, totalling 613 microbiome samples and 628 cytokine samples. Microbial taxa that appeared in less than 5% of the samples were removed, leaving 213 taxa. Zeros were imputed by 0.5 before centered log-ratio transformation (CLR) was applied. Cytokines that were detected in less than 10% of the samples (FGF basic and IL-17) were removed, leaving 27 cytokines. Zeros were imputed using half of the smallest observed positive values for each cytokine, and log2 transformation was applied before being analyzed by multiTEMPTED. multiTEMPTED used a kernel ridge penalty of 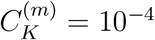 for all modalities.

#### B.2 iPOP Exercise Study

Data were downloaded from http://hmp2-data.stanford.edu/. The initial feature set comprised 728 metabolites (LC-MS), 710 lipid species (Lipidyzer), 109 targeted protein markers (immunoassays), and 260 untargeted protein markers (SWATH-MS). Fifty-four metabolites with low prevalence across samples were excluded, retaining 674. Targeted and untargeted protein markers were merged into a single protein modality of 369 features. All retained features were log10-transformed and then standardized to zero mean and unit variance across subjects and time points.

Because individual lipid species within the same biochemical class are highly correlated, we reduced the lipid modality by grouping lipids into 13 classes: Cholesteryl Ester (CE), Ceramide (CER), Diacylglycerol (DAG), Dihydroceramide (DCER), Free Fatty Acid (FFA), Hexosylceramide (HCER), Lactosylceramide (LCER), Lysophosphatidylcholine (LPC), Lysophosphatidylethanolamine (LPE), Phosphatidylcholine (PC), Phosphatidylethanolamine (PE), Sphingomyelin (SM), and Triacylglycerol (TAG). Within the TAG class, lipids were further divided by chain length into mid-chain (MCFA, ≤ 12 carbons), long-chain (LCFA, 13–21 carbons, subgrouped by degree of unsaturation), and very-long-chain (VLCFA, ≥ 22 carbons) categories. Within each class or TAG subclass, PCA was applied and the components collectively explaining at least 95% of within-class variance were retained as summary features, labeled by class and index (e.g., `CE.pc.1` for the first principal component of the cholesteryl ester class). This aggregation yielded 122 lipid summary features, which were standardized in the same manner as the other modalities. For both multiTEMPTED and MEFISTO, *r* = 3 components were fitted with equal modality weights. multiTEMPTED used a kernel ridge penalty of 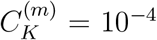 for all modalities.

### C Simulation Details

#### Common setup

Unless stated otherwise, synthetic data are generated from model (1) with *M* = 3 modalities and *r* = 3 components. Independent *N* (0, *σ*^2^) noise is added to each 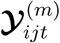, and modality scales 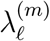 decay across components so that components are ordered by signal strength. Temporal loadings 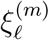 are drawn from different analytic shapes (monotone, single-bend, and multi-bend; see Figure 3), normalized to unit *L*_2_ norm on *T* = [0, 1]. Estimation uses *r* = 3, no transformation or centering, and kernel ridge penalty 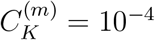.

In Simulations 2 and 3, MEFISTO is run with groups defined as subjects and time as the continuous covariate, with 1,000 iterations, the “`slow`” convergence criterion, and Gaussianprocess hyperparameter optimization enabled (its most accurate configuration). Neither method takes the input of the subjects’ group labels. Because a MEFISTO factor supplies one value per (subject, time) sample rather than a separated subject loading and temporal curve, we recover the two by taking the rank-one SVD of each factor’s subject-by-time-bin matrix: the left singular vector plays the role of *a*_*ℓ*_ and the right singular vector that of *ξ*_*ℓ*_.

**Simulation 1 setup** *n* = 30 subjects, *p*^(*m*)^ = 20 features per modality, *q*_*im*_ = 8 sampling times per subject and modality drawn independently and uniformly on [0, 1], so no two subjects or modalities share a time point. Every (modality, component) pair receives a distinct temporal shape. Noise is varied over *σ* ∈ {0.1, 0.5, 1.0, 2.0, 4.0} with 20 random seeds at each level. Only multiTEMPTED is run in this scenario.

**Simulation 2 setup**. *n* = 12 subjects, *M* = 3 modalities, *p*^(*m*)^ = 24 features, *r* = 3, *σ* = 0.1, over 10 random seeds. We did not run more simulation runs due to the high time cost of running MEFISTO in this setup. Each (modality, component) pair is assigned its own temporal shape (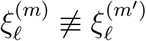 for *m ?*= *m*^*′*^), and each subject–modality combination was sampled at time points independently drawn from a uniform distribution.

**Simulation 3 setup**. *n* = 15 subjects, *M* = 3 modalities, *p*^(*m*)^ = 24 features, *r* = 3, *σ* = 1, over 100 random seeds. Temporal loadings are *shared* across modalities: 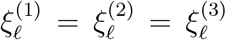. Subjects are partitioned into *r* = 3 groups and features into *r* matching subsets per modality, so component *ℓ* is active only for group-*ℓ* subjects and subset-*ℓ* features, with small off-block values. Sampling follows a fixed schedule of 8 equally spaced nominal time points, jittered per subject by *N*(0, 0.02^2^). All modalities are co-measured at each subject’s realized visit time. This mirrors the structure of clinical cohort studies, where one visit supplies all assays, and visits could drift from the nominal schedule. This setup also avoids a degenerate regime for MEFISTO in which identical covariate values across subjects trigger an expensive group-by-group covariance optimization that makes a hundred-seed run infeasible.

